# AncestralClust: Clustering of Divergent Nucleotide Sequences by Ancestral Sequence Reconstruction using Phylogenetic Trees

**DOI:** 10.1101/2021.01.08.426008

**Authors:** Lenore Pipes, Rasmus Nielsen

## Abstract

**Motivation:** Clustering is a fundamental task in the analysis of nucleotide sequences. Despite the exponential increase in the size of sequence databases of homologous genes, few methods exist to cluster divergent sequences. Traditional clustering methods have mostly focused on optimizing high speed clustering of highly similar sequences. We develop a phylogenetic clustering method which infers ancestral sequences for a set of initial clusters and then uses a greedy algorithm to cluster sequences.

**Results:** We describe a clustering program *AncestralClust*, which is developed for clustering divergent sequences. We compare this method with other state-of-the-art clustering methods using datasets of homologous sequences from different species. We show that, in divergent datasets, AncestralClust has higher accuracy and more even cluster sizes than current popular methods.

**Availability and implementation:** AncestralClust is an Open Source program available at https://github.com/lpipes/ancestralclust

**Supplementary information:** Supplementary figures and table are available online.

## 1 Introduction

Traditional clustering methods such as UCLUST (Edgar, 2010) and CD-HIT (Fu *et al*., 2012) use hierarchical or greedy algorithms that rely on user input of a sequence identity threshold. These methods were developed for high speed clustering of a high quantity of highly similar sequences (Ghodsi *et al*., 2011; Li *et al*., 2001; Edgar, 2010) and, generally, these methods are considered unreliable for identity thresholds <75% because of either the poor quality of alignments at low identities (Zou *et al*., 2018) or because the performance of the method drops dramatically with low identities (Huang *et al*., 2010). At low identities, these methods produce uneven clusters where the majority of sequences are contained in only one or a few clusters (Chen *et al*., 2018). A high variance in cluster sizes may reduce the utility of clustering for many practical purposes since the goal of clustering typically is to reduce computational complexity of downstream analyses that are limited by the size of the largest clusters. Clustering of divergent sequences is an important step in genomics analysis because it allows for an early divide-and-conquer strategy that will significantly increase the speed of downstream analyses (Zheng *et al*., 2018) and many fundamental questions in metagenomics can be addressed by clustering of divergent sequences, such as the identification of gene families and identification of sequences at the order, class, or phylum taxonomic levels. Currently, there are no clustering methods that can accurately cluster large taxonomically divergent metabarcoding reference databases such as the Barcode of Life database (Ratnasingham and Hebert, 2007) in relatively even clusters. Only a few other methods, such as SpClust (Matar *et al*., 2019) and TreeCluster (Balaban *et al*., 2019), exist for clustering potentially divergent sequences. SpClust creates clusters based on the use of Laplacian Eigenmaps and a Gaussian Mixture Model based on a similarity matrix calculated on all input sequences. While this approach is highly accurate, the calculation of an all-to-all similarity matrix is a computationally demanding. For example, an all-by-all comparison for clustering 8 million environmental DNA reads by Rusch *et al*. (2007) took >1 year on a 100-CPU cluster. TreeCluster uses user-specified constraints for splitting a phylogenetic tree into clusters. However, TreeCluster requires an input tree and even though some phylogenetic methods exist to estimate trees from a large number of sequences (Stamatakis, 2014), it can also be prohibitively slow for large numbers of divergent sequences where a phylogenetic tree is difficult to estimate reliably. With the increasing size of reference databases (Schoch *et al*., 2020), there is a need for new computationally efficient methods that can cluster divergent sequences. Here we present AncestralClust which is specifically developed for clustering of divergent metabarcoding reference sequences in clusters of relatively even size.

## 2 Methods

To cluster divergent sequences, we developed AncestralClust written in C (Figure 1). Firstly, *r* random sequences are chosen and the sequences are aligned pairwise using the wavefront algorithm (Marco-Sola *et al*., 2020). A Jukes-Cantor distance matrix is constructed from the alignments and a neighbor-joining phylogenetic tree is constructed. The Jukes-Cantor model is chosen for computational speed, but more complex models could in principle be used to potentially increase accuracy but also increase computational time. The *k* − 1 longest branches in the tree are then cut to yield *k* clusters. These subtrees comprise the initial starting clusters. The sequences in each starting cluster are aligned in a multiple sequence alignment using kalign3 (Lassmann, 2020), a neighbor joining tree is constructed from each starting cluster, and midpoint rooted. The ancestral sequences at the root of the tree of each cluster is estimated using the maximum of the posterior probability of each nucleotide using standard programming algorithms from phylogenetics (see e.g., Yang, 2014). The ancestral sequences are used as the representative sequence for each cluster. Next, the rest of the sequences are assigned to each cluster based on the shortest nucleotide distance from the wavefront alignment between the sequence and the *k* ancestral sequences. If the shortest distance to any of the *k* ancestral sequences is larger than the average distance between clusters, the sequence is saved for the next iteration. We iterate this process until all sequences are assigned to a cluster. In each iteration after the first iteration, a cut of a branch in the phylogenetic tree is chosen if the branch is longer that the average length of branches cut in the first iteration. In praxis, only one or two iterations are needed for most data sets if *r* is defined to be sufficiently large.

**Figure 1.**
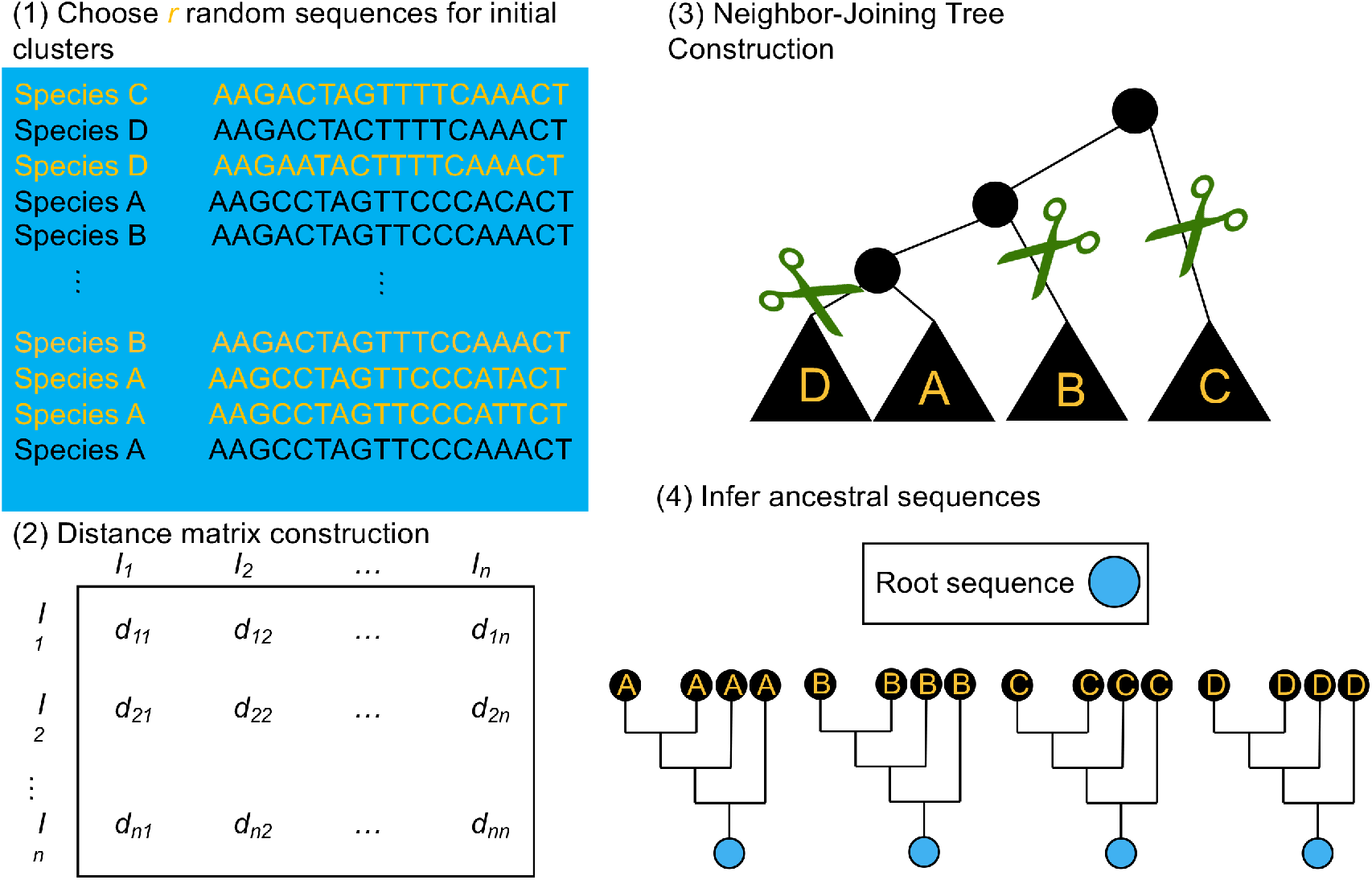
Overview of AncestralClust. In (1), *r* random sequences are chosen for the initial clusters. (2) Using the *r* sequences a distance matrix is constructed. Using the distance matrix, a neighbor-joining tree is constructed and *k* − 1 cuts are made to create *k* clusters. In (4), each cluster is multiple-sequence aligned and the ancestral sequences are reconstructed in the root node of each tree. The rest of the unassigned sequences are then aligned to the ancestral sequences of each cluster and the shortest distance to each ancestral sequence is calculated. The process is iterated until all sequences are assigned to a cluster.

We compared AncestralClust to three other state-of-the-art clustering methods: UCLUST (Edgar, 2010), CD-HIT (Fu *et al*., 2012), and SpClust (Matar *et al*., 2019). We used a variety of measurements to assess the accuracy and evennness of the clustering. We calculated two traditional measures of accuracy, purity and normalized mutual information (NMI), used in Bonder *et al*. (2012). The purity of clusters is calculated as:

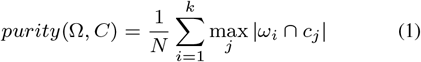

where Ω = {*w*_1_, *w*_2_, …, *w*_*k*_} is the set of *k* clusters, *C* = {*c*_1_, *c*_2_, …, *c*_*d*_} is the set of *d* taxonomic groups, and *N* is the total number of sequences. NMI is calculated as:

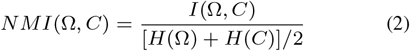

where mutual information gain is *I*(Ω, *C*) and *H* is the entropy function. To measure the evenness of the clusters, we used the coefficient of variation, which is calculated as:

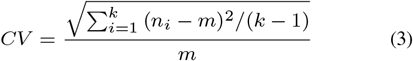

where *n*_*i*_ = |*ω*_*i*_| is the number of sequences in cluster *i* and 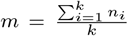 is the mean size of the clusters. We also used a taxonomic incompatibility measure to assess the accuracy of the clusters. A taxonomic incompatibility is assigned if two taxonomic groups both exist in two different clusters. One taxonomic group can be split into multiple clusters, but if taxonomic groups are monophyletic, two groups should not both be found in more than one cluster. The total taxonomic incompatibility is then calculated by summing over all species found in the data set. More precisely, let *S*_*ω*_(*c*_*i*_, *c*_*j*_) be an indicator variable that returns one if species *c*_*i*_ and *c*_*j*_ are both found in cluster *ω* and zero otherwise, then the taxonomic incompatibility is defined as:

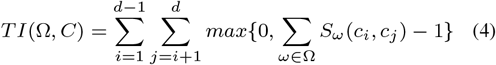

Notice that this measure does not penalize paraphyletic clusters but only penalizes clustering that is strictly incompatible with a phylogentic tree, i.e. polyphyletic clusters.

All three measures are very sensitive to both the number of clusters and the variance in cluster size. For example, if all sequences are assigned to different clusters or if all sequences are assigned to the same cluster, the taxonomic incompatibility is per definition zero. In general, taxonomic incompatibility has the potential to be highest, and NMI has the potential to be lowest, when there is an intermediate amount of clusters of equal size. With high variance in cluster size there is less potential for generating clusters with high phylogentic incompatibility of low NMI. To address this issues in order to allow fair comparison when numbers of clusters and variance in cluster sizes vary, we calculated the *relative purity, relative NMI*, and *relative incompatibility*. These measures were calculated by scaling them relative to their expected values under random assignments given the number of clusters and the cluster sizes. We estimated *relative NMI* by dividing the raw NMI score by the average NMI of 10 clusterings, in which sequences have been assigned at random with equal probability to clusters, such that the cluster sizes are the same as the cluster sizes produced in the original clustering. The same procedure was used to convert the purity measure into *relative purity* and the taxonomic incompatibility measure into *relative incompatibility*.

## 3 Results

To first assess performance of clustering methods on divergent nucleotide sequences, we used 100 random samples of 10,000 sequences from three metabarcode reference databases (16, 18S, and Cytochrome Oxidase I (COI)) from the CALeDNA project Curd *et al*. (2019). We first compare our method on this dataset against UCLUST and CD-HIT, which are arguably the most popular clustering methods. We were unable to perform clustering of this dataset using SpClust because the program did not complete in a feasible amount of time (we allowed for a week of computational time to complete the clusterings of 100 random samples for each method). In the main figures, we display the results against UCLUST but because of the high number of clusters and high coefficient of variation values of CD-HIT (see Figures S2, S7 and S8) we display the CD-HIT results in the supplement. For CD-HIT, we used the lowest possible similarity threshold, which is 80%, to attempt to create clusters of similar sizes to AncestralClust.

We first compared AncestralClust against UCLUST using *relative NMI* and the Coefficient of Variation for species (Figure 2), genus, family, order, class, and phylum levels (Figure S1) across 16S, 18S, and COI metabarcodes. We used *r* = 300 random initial sequences, which is 3% of the total number of sequences in each sample and *k* = 15 initial clusters. Results for CD-HIT (Figure S2) show that CD-HIT creates hundreds of clusters for every barcode with a high Coefficient of Variation and tends to have a lower *relative NMI* than AncestralClust. Notice that the relative NMI tends to be higher with a lower coefficient of variation for AncestralClust across all barcodes. This suggests, that for these divergent eDNA sequences, AncestralClust provides clusterings that are more even in size and that are more consistent with conventional taxonomic assignment. As a second and third measure of accuracy we measured *relative purity* and *relative incompatibility* and coefficient of variation using AncestralClust, UCLUST, and CD-HIT for the same datasets under the same running conditions. Notice in Figures 3 and 4, AncestralClust tends to create balanced clusters with higher *relative purities* and lower *relative taxonomic incompatibilities* compared to UCLUST at all taxonomic levels. Similar results are seen for metabarcode 18S *relative incompatibility* (Figure S3), although at the species level the two methods now perform similarly. However, for *relative purity* for 18S, the two methods perform similarly (Figure S4). For *relative incompatibility* for metabarcode 16S (Figure S5), AncestralClust performs noticeably better than UCLUST at the species and genus levels but at the family, order, class, and phylum levels it has either the same or slightly more taxonomic incompatibility, but substantially lower Coefficient of Variation of the cluster sizes. Also, at the species, genus, and family levels, there is a clear negative correlation between UCLUST *relative incompatibility* and Coefficient of Variation. This illustrates the observation that clusterings with a higher variance in cluster sizes tend to generate lower taxonomic incompatibility. For *relative purity* for 16S, AncestralClust has noticeably higher *relative purities* than UCLUST at every taxonomic level (Figure S6). For CD-HIT, *relative purities* of 16S, 18S, and COI tends to be similar or lower than AncestralClust at every taxonomic level (Figure S7. Additionally, *relative incompatibility* of 16S and 18S tends to be similar or higher than AncestralClust at every taxonomic level (Figure S8). For COI, *relative incompatibility* tends to be higher or similar at species, genus, and family levels, but lower at order, class, and phylum levels, but with substantially higher coefficient of variation in cluster size.

**Figure 2.**
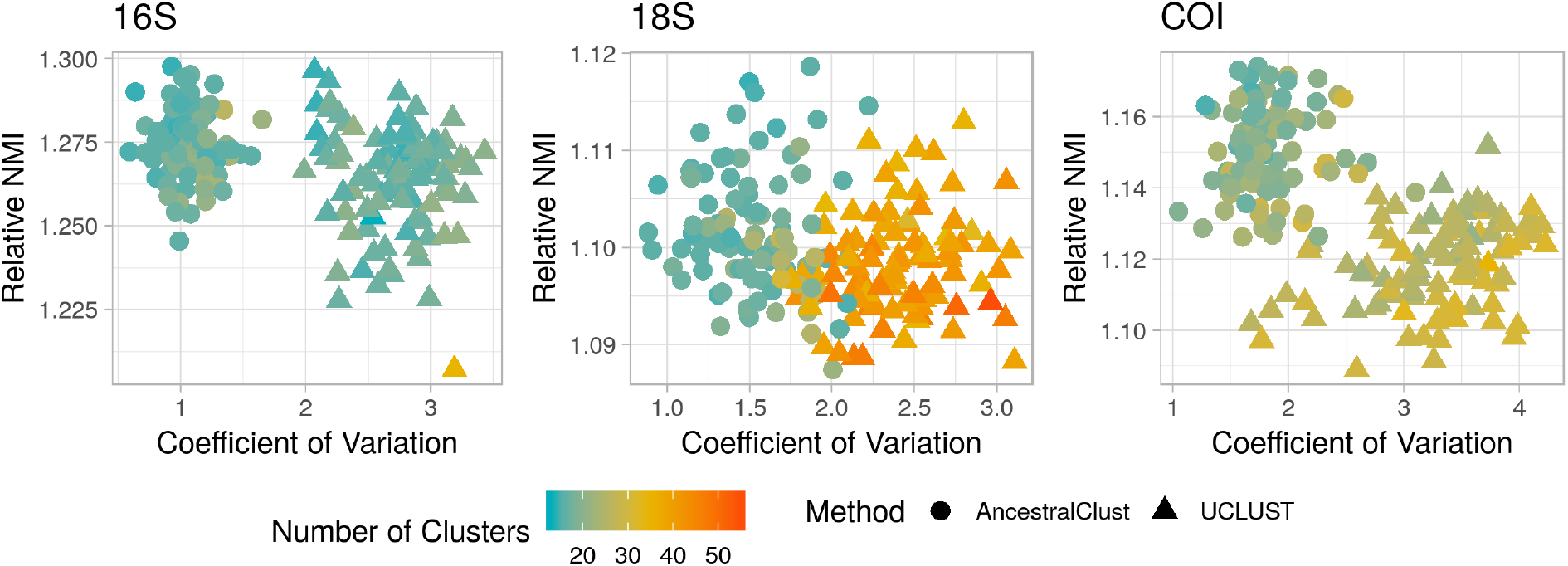
Relative NMI at the family level against coefficient of variation for AncestralClust and UCLUST for 100 samples of 10,000 randomly chosen 16S, 18S, and COI reference sequences from the CALeDNA Project (Curd et al., 2019). The similarity threshold for UCLUST was 0.58. For AncestralClust, we used 300 initial random sequences with 15 initial clusters. Relative NMI was calculated by dividing NMI by the average of 10 random samples of the same fixed cluster size.

**Figure 3.**
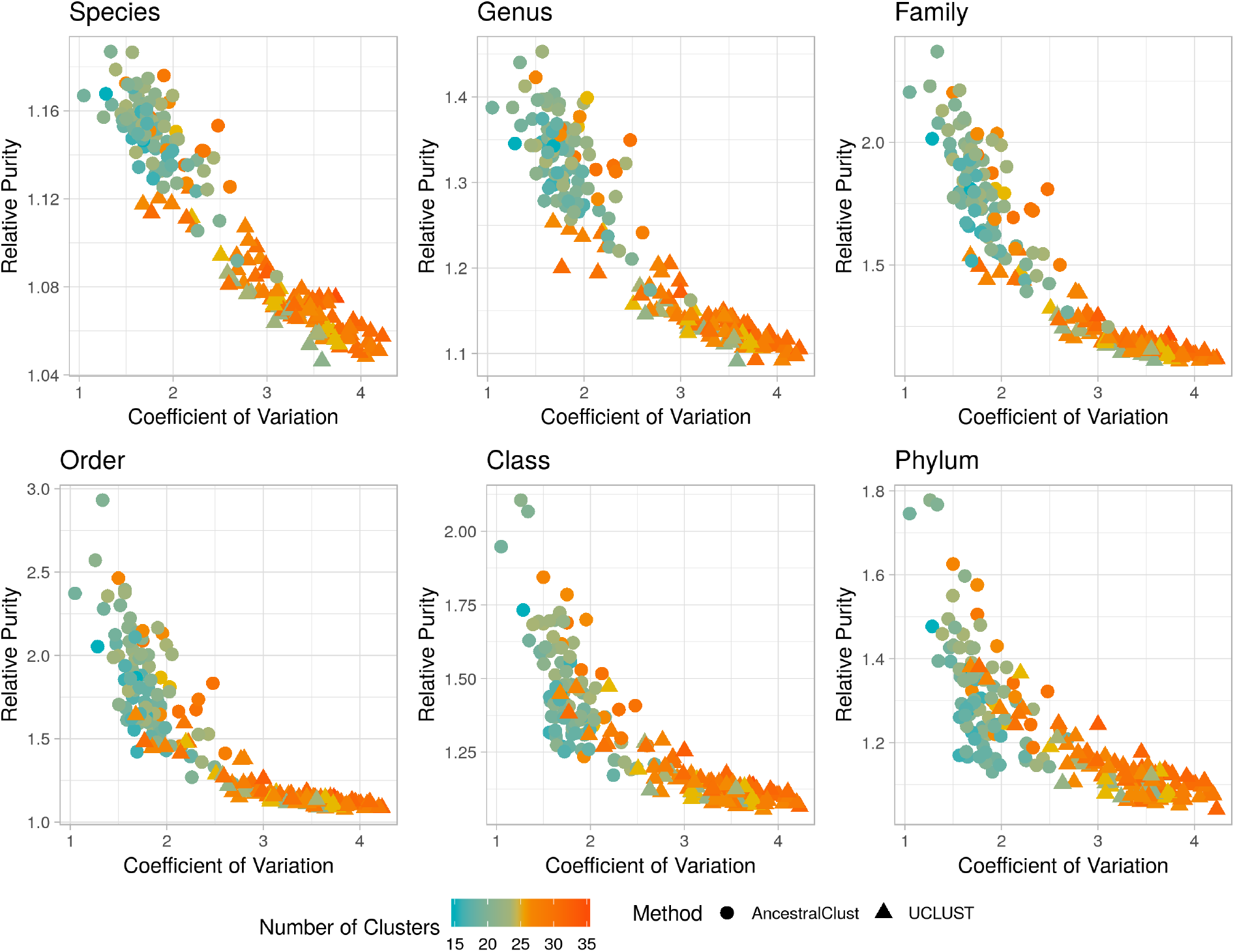
Relative purity against coefficient of variation for AncestralClust and UCLUST for 100 samples of 10,000 randomly chosen COI reference sequences. COI reference sequences are from the CALeDNA Project (Curd et al., 2019). The similarity threshold for UCLUST was 0.58. For AncestralClust, we used 300 initial random sequences with 15 initial clusters.

**Figure 4.**
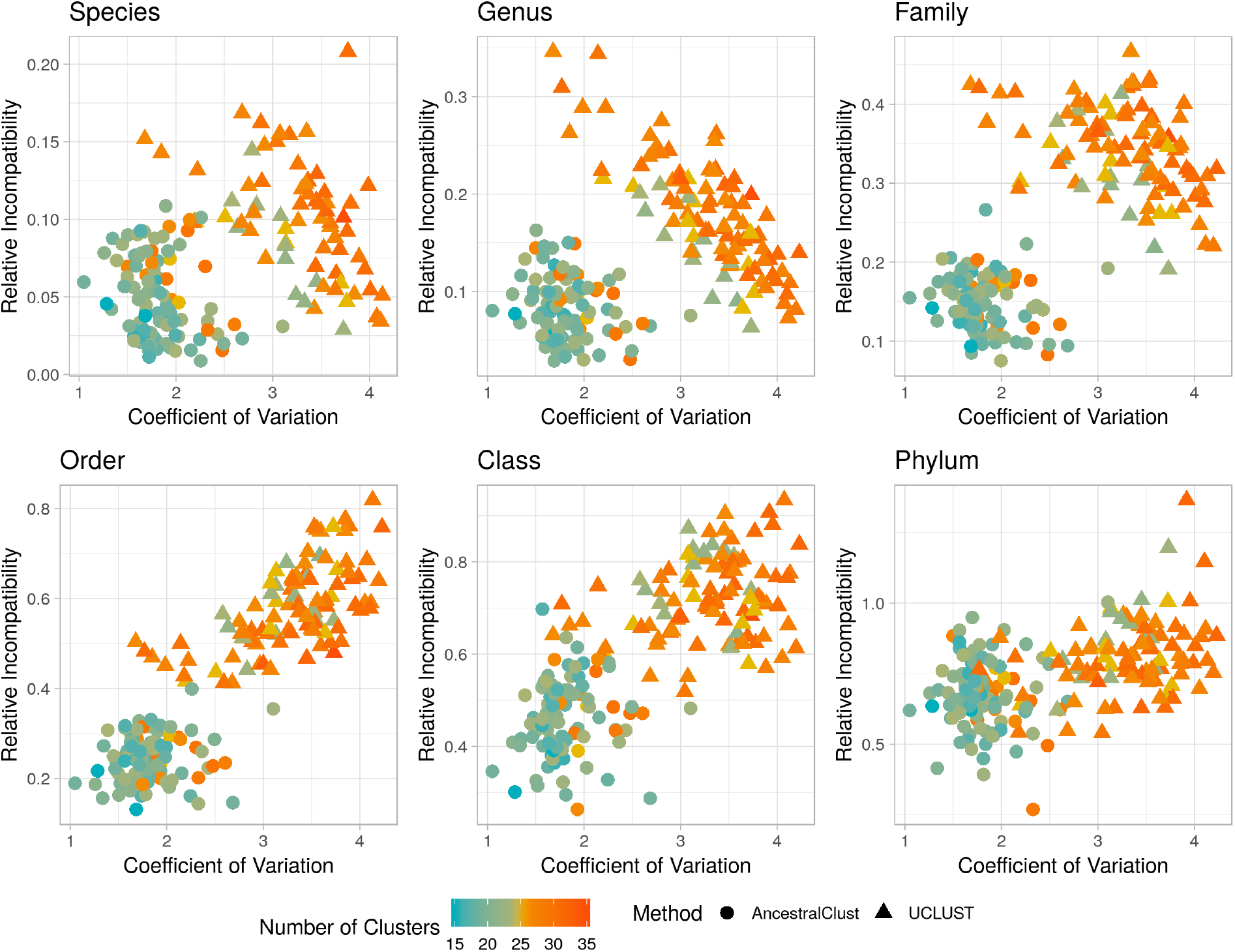
Relative incompatibility against coefficient of variation for AncestralClust and UCLUST for 100 samples of 10,000 randomly chosen COI reference sequences. COI reference sequences are from the CALeDNA Project (Curd et al., 2019). The similarity threshold for UCLUST was 0.58. For AncestralClust, we used 300 initial random sequences with 15 initial clusters.

Next, we analyzed two datasets with different properties: one dataset of diverse species from the same gene and another dataset of 6 paralogous genes from species of the same phylum. In the first dataset, we expect the sequences to cluster according to species. In the second dataset, we expect the sequences to cluster according to genes. We compared AncestralClust to two commonly used clustering programs (UCLUST and CD-HIT) and one clustering program designed for divergent sequences, SpClust. The first dataset contained 13,043 sequences from the COI CaleDNA database from 11 divergent species that were from 7 different phyla and 11 different classes and the second data set contained sequences from 6 different genes from taxonomically similar species. First, we compared all methods using 13,043 COI sequences from the 11 different species (Table 1). Ideally, we expect these sequences to form 11 different clusters, each including all the sequences from one species. We chose identity thresholds to enforce the expected number of clusters for each method. We were unable to form 11 clusters using CD-HIT because the program does not allow clustering of sequences with identity thresholds < 80% at default parameters. For SpClust, we used the three precision modes available for the method. In this analysis, AncestralClust achieved a perfect clustering (the raw purity was 1 and *relative incompatibility* was 0) and it had the second lowest memory usage, although it was the second slowest. UCLUST was one of the fastest methods and used the least amount of memory but had the second lowest purity with third highest *relative NMI* values. SpClust only identified one cluster, with a computational time of ∼2 days. In comparison, AncestralClust took ∼5 minutes and UCLUST used < 1 second.

**Table 1.** Comparisons of clustering methods using 13,043 COI sequences from 11 different species. The list of species can be found in Table S1. Incompatibility was calculated at the taxonomic rank of species. For UCLUST, the identity thresholds were chosen to force the expected 11 number of clusters. For CD-HIT, the lowest possible identity was chosen which is 0.8. In the case of SpClust, Coefficient of Variation cannot be calculated for 1 cluster. SpClust clusters were created with version 2.

| Method | # of clusters | Time (sec) | Mem (MB) | Relative Purity (species) | Relative Incompat. (species) | Relative NMI (species) | Coeff. of Var. |
| --- | --- | --- | --- | --- | --- | --- | --- |
| AncestralClust | 11 | 293.2 | 19.3 | 3.6271 | 0 | 551.09 | 0.8574 |
| UCLUST | 11 | <1 | 9.9 | 3.1201 | 0.0182 | 474.63 | 0.8300 |
| CD-HIT | 24 | 5.86 | 43.9 | 4.1234 | 0 | 241.66 | 1.2031 |
| SpClust (fast) | 1 | 152046.5 | 2678.9 | 1 | 0 | 1 | - |
| SpClust (moderate) | 1 | 188172.9 | 6457.6 | 1 | 0 | 1 | - |
| SpClust (maxPrecision) | 1 | 189577.1 | 6452.5 | 1 | 0 | 1 | - |

Next, we analyzed ‘genomic set 1’ from Matar *et al*. (2019), which consists of 39 sequences from 6 homologous genes (FCER1G, S100A1, S100A6, S100A8, S100A12, and SH3BGRL3 in Table 2). We expect these sequences to form 6 clusters. We varied the identity thresholds for UCLUST using thresholds 0.4, 0.6, and 0.8. For CD-HIT, we used the lowest identity threshold available on default parameters which is 0.8. Since this dataset contained 6 different genes, we calculated *relative NMI* using genes as the classes and did not use incompatibility as an accuracy measure. Only AncestralClust and UCLUST produced the expected number of clusters, and among the methods and parameters that created the expected number of clusters, AncestralClust had the highest *relative purity* value. AncestralClust was the second slowest method and had the highest memory requirements due to the wavefront algorithm alignment which is *𝒪* (*s*^2^) in memory requirements, where *s* is the alignment score. Since alignments were performed using 6 different genes that were longer than 1.5kb (the average sequence length was 2,387.9bp and the longest sequence was 5,363bp), this resulted in a high value of *s*. SpClust had the highest *relative NMI* and lower *relative purity* than AncestralClust for all precision modes, however, it failed to produce the expected number of clusters and found fewer clusters with a higher Coefficient of Variation than AncestralClust, making the results difficult to compare.

**Table 2.** Comparisons of clustering methods using 39 sequences from 6 homologous genes from Matar et al. (2019).’id’ refers to the identity threshold used. We used identity thresholds of 0.4, 0.6, and 0.8 for UCLUST. We used precision levels of fast, moderate, and maximum for SpClust using version 1 since version 2 only produced 1 cluster for all modes.

| Method | # of clusters | Time (sec) | Memory (Mb) | Relative Purity | Relative NMI | Coeff. of Var. |
| --- | --- | --- | --- | --- | --- | --- |
| AncestralClust | 6 | 370.3 | 412.0 | 1.7619 | 1.8660 | 0.3982 |
| UCLUST (id=0.4) | 6 | 1 | 15.4 | 1.3810 | 1.5667 | 0.5396 |
| UCLUST (id=0.6) | 19 | 1 | 20.1 | 2.1538 | 1.4379 | 0.7166 |
| UCLUST (id=0.8) | 29 | 1.9 | 20.4 | 1.6923 | 1.1717 | 0.4565 |
| CD-HIT (id=0.8) | 31 | 0.48 | 39.9 | 1.2308 | 1.0950 | 0.4574 |
| SpClust (fast) | 4 | 44.6 | 166.2 | 1.4167 | 2.2463 | 0.8432 |
| SpClust (moderate) | 4 | 112.5 | 166.1 | 1.6818 | 2.4335 | 0.6453 |
| SpClust (max precision) | 4 | 570.1 | 166.0 | 1.6818 | 2.9449 | 0.6809 |

## 4 Conclusions

We developed a phylogenetic-based clustering method, AncestralClust, specifically to cluster divergent metabarcode sequences. We performed a comparative study between AncestralClust and widely used clustering programs UCLUST and CD-HIT, and for divergent sequences, SpClust. UCLUST is substantially faster than AncestralClust and should be the preferred method if computational speed is the main concern. However, AncestralClust tends to form clusters of more even size with lower *relative taxonomic incompatibility* and higher *relative NMI* and *relative purity* than other methods, for the relatively divergent sequences analyzed here. We recommend the use of AncestralClust when sequences are divergent, especially if a relatively even clustering is also desirable, for example for various divide-and-conquer approaches where computational speed of downstream analyses increases faster than linearly with cluster size.

