## Supplementary Material for "AncestralClust: Clustering of Divergent Nucleotide Sequences by Ancestral Sequence Reconstruction using Phylogenetic Trees"

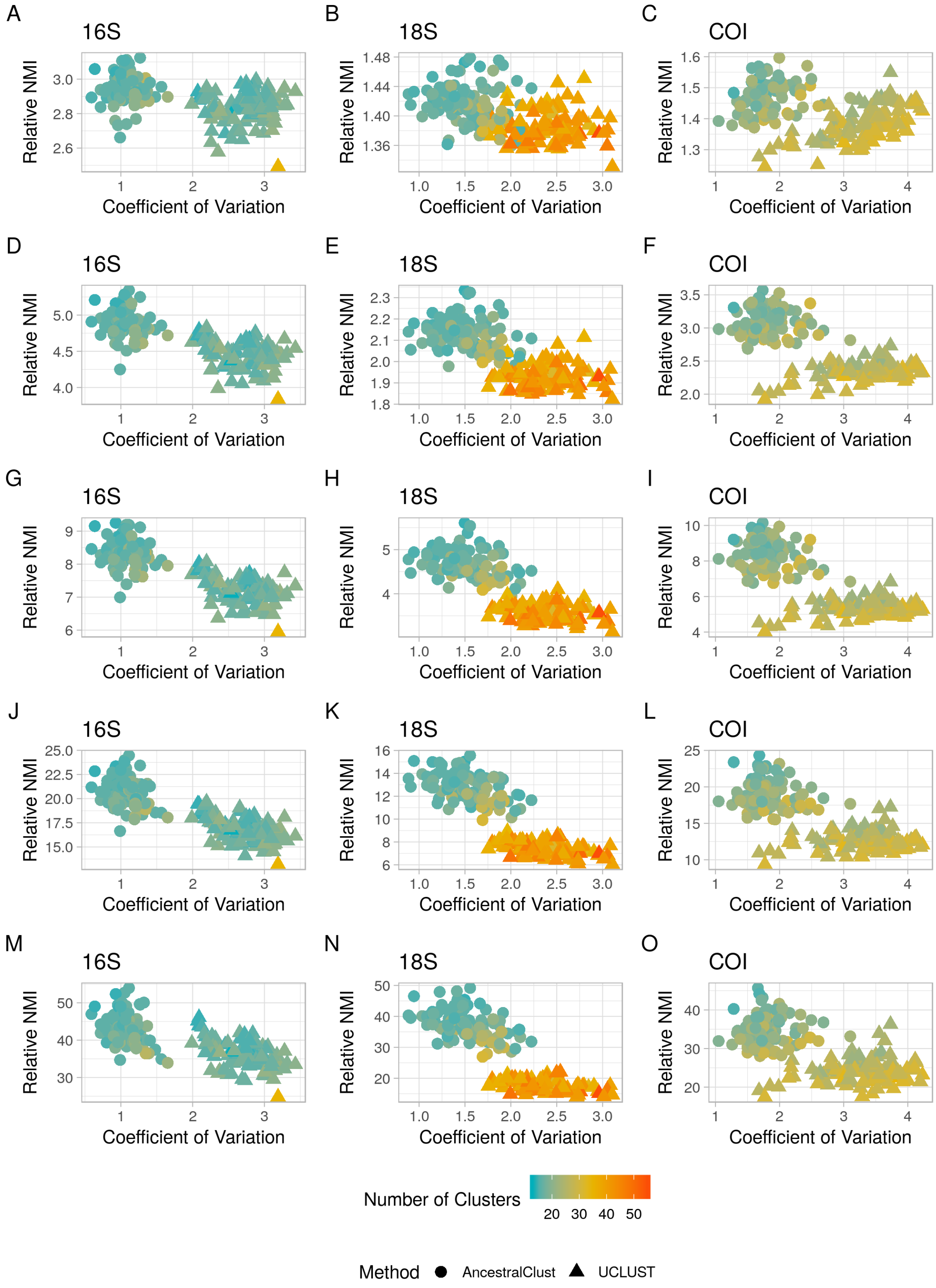


**Figure S1.** Relative NMI against coefficient of variation for AncestralClust and UCLUST for 100 samples of 10,000 randomly chosen 16S, 18S, and COI reference sequences for taxonomic levels genus (A-C), family (D-F), order (G-I), class (J-L), and phylum (M-O). All reference sequences are from the CALeDNA Project (Curd et al., 2019). The similarity threshold for UCLUST for 16S and 18S is 0.58, and for COI the similarity threshold is 0.62. For AncestralClust, we used 300 initial random sequences with 15 initial clusters.

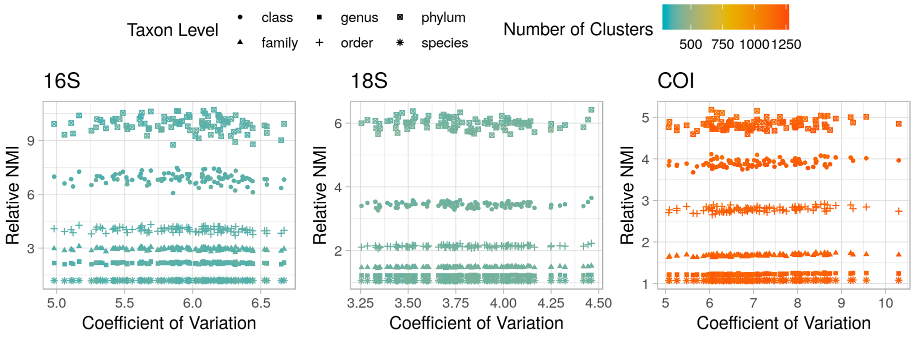


**Figure S2.** Relative NMI at all taxon levels for CD-HIT against coefficient of variation for 100 samples of 10,000 randomly chosen 16S, 18S, and COI reference sequences from the CALeDNA Project (Curd et al., 2019). The similarity threshold for CD-HIT is 0.8. Relative NMI was calculated by dividing NMI by the average of 10 random samples of the same fixed cluster size.

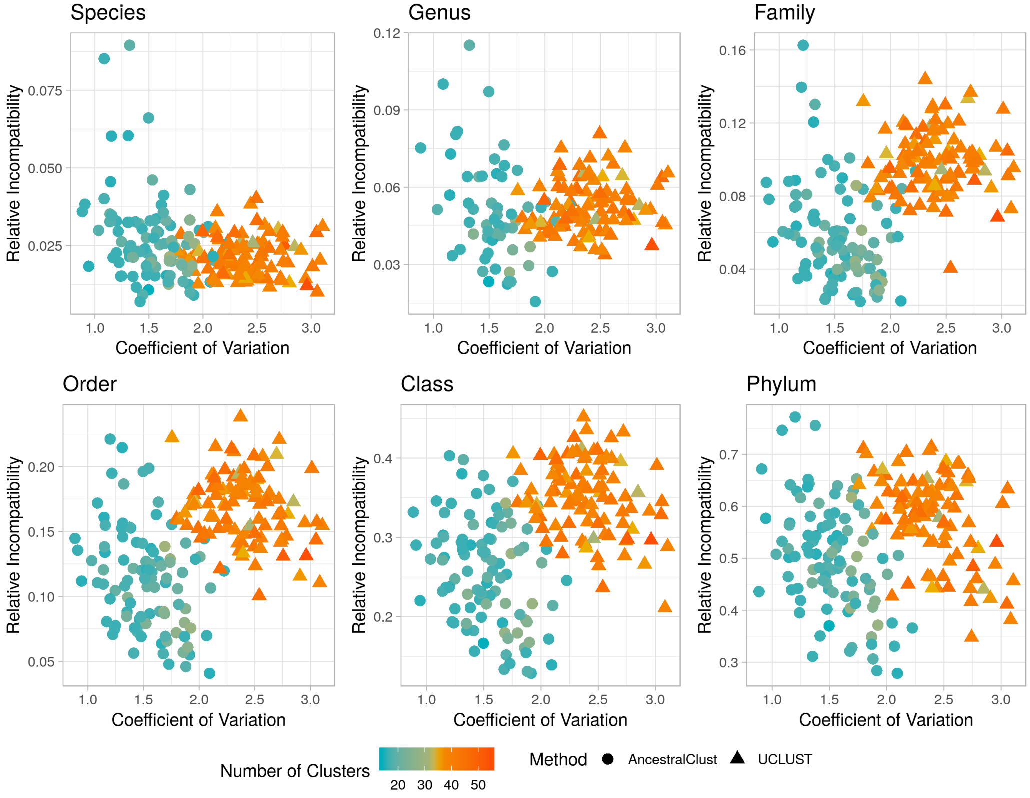


**Figure S3.** Relative incompatibility against coefficient of variation for AncestralClust and UCLUST for 100 samples of 10,000 randomly chosen 18S reference sequences. 18S reference sequences are from the CALeDNA Project (Curd et al., 2019). The similarity threshold for UCLUST was 0.58. For AncestralClust, we used 300 initial random sequences with 15 initial clusters.

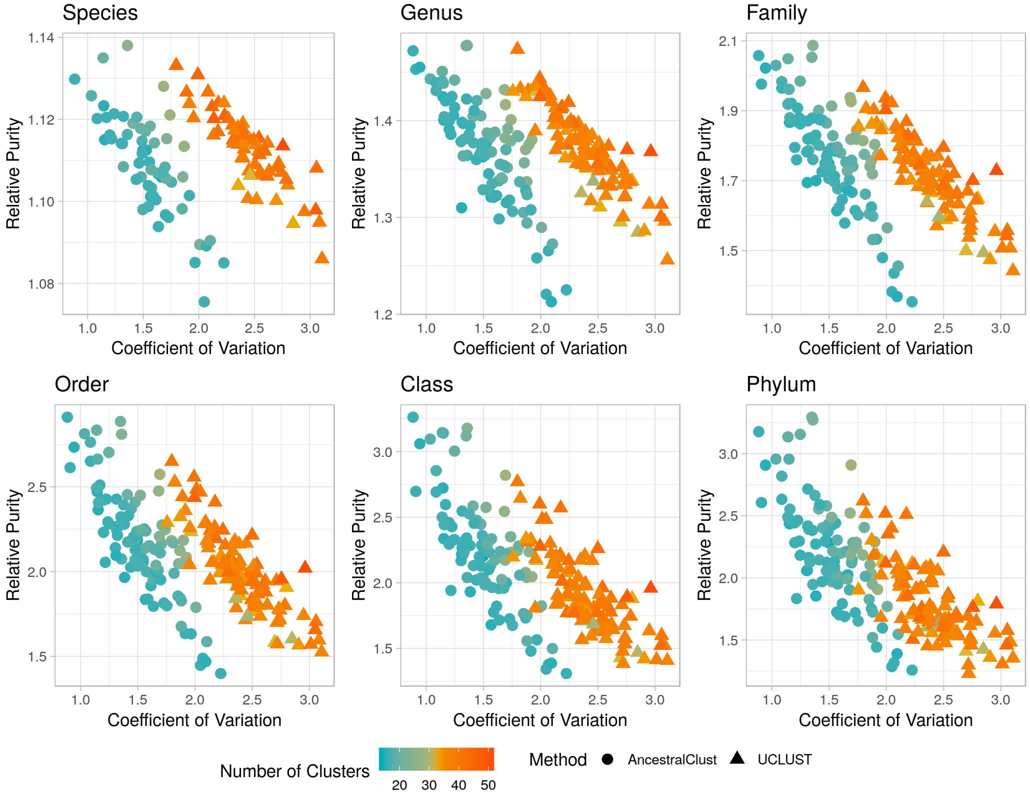


**Figure S4.** Relative purity against coefficient of variation for AncestralClust and UCLUST for 100 samples of 10,000 randomly chosen 16S reference sequences. 18S reference sequences are from the CALeDNA Project (Curd et al., 2019). The similarity threshold for UCLUST was 0.58. For AncestralClust, we used 300 initial random sequences with 15 initial clusters.

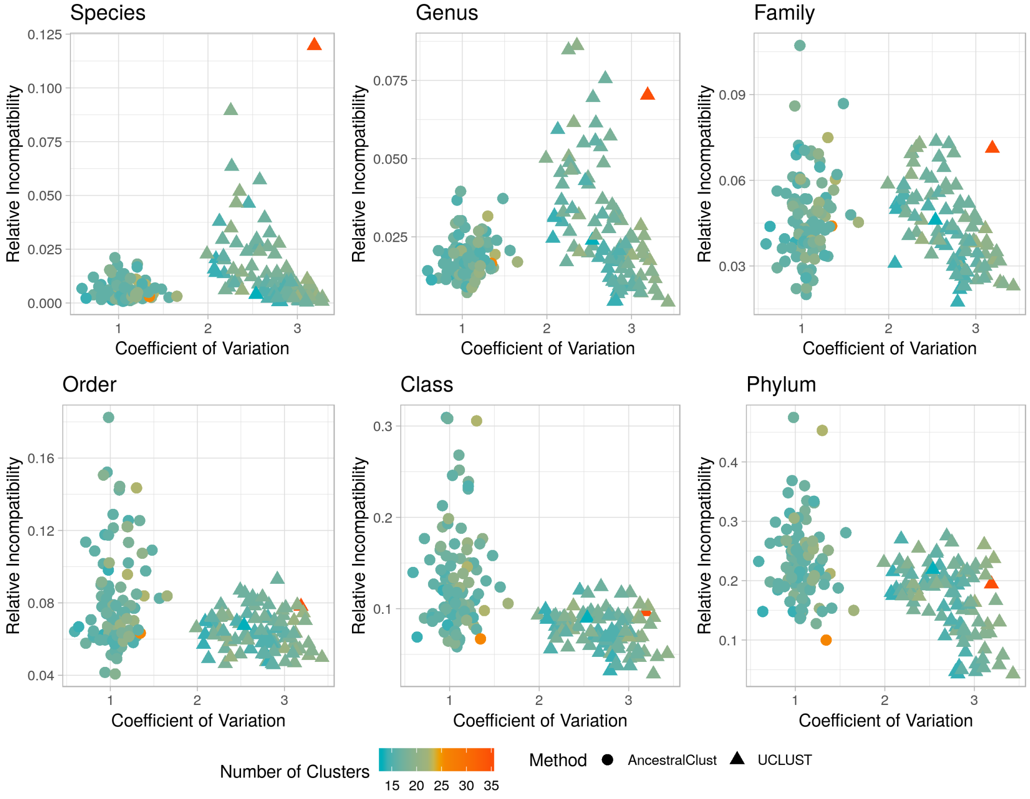


**Figure S5.** Relative incompatibility against coefficient of variation for AncestralClust and UCLUST for 100 samples of 10,000 randomly chosen 16S reference sequences. 16S reference sequences are from the CALeDNA Project (Curd et al., 2019). The similarity threshold for UCLUST was 0.58. For AncestralClust, we used 300 initial random sequences with 15 initial clusters.

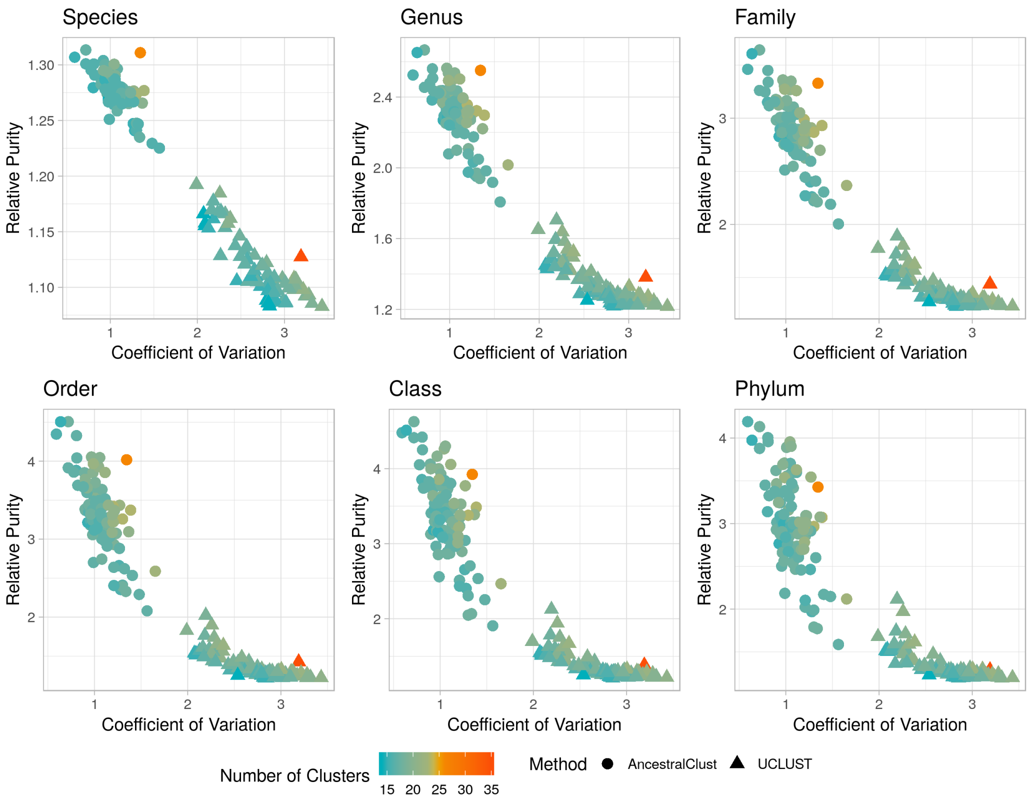


**Figure S6.** Relative purity against coefficient of variation for AncestralClust and UCLUST for 100 samples of 10,000 randomly chosen 16S reference sequences. 16S reference sequences are from the CALeDNA Project (Curd et al., 2019). The similarity threshold for UCLUST was 0.58. For AncestralClust, we used 300 initial random sequences with 15 initial clusters.

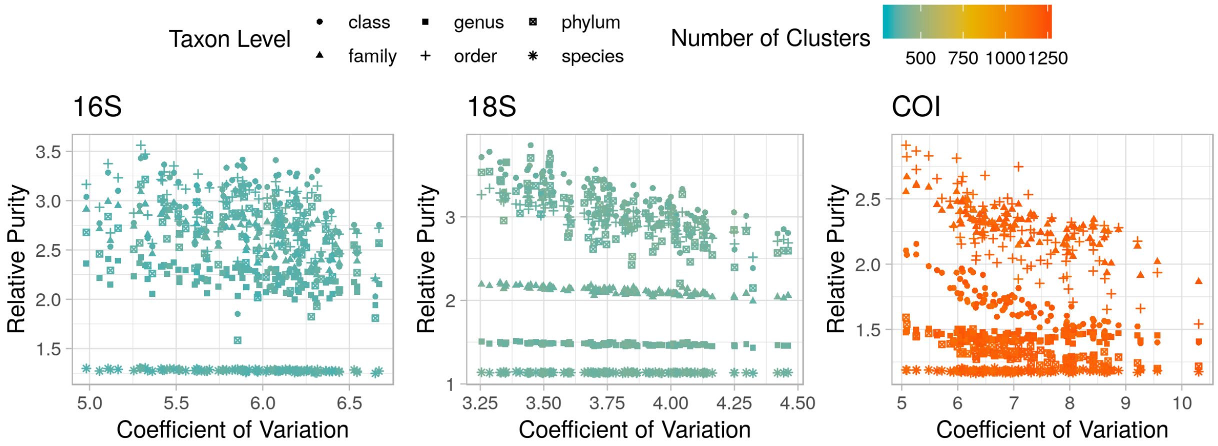


**Figure S7.** Relative purity at all taxon levels for CD-HIT against coefficient of variation for 100 samples of 10,000 randomly chosen 16S, 18S, and COI reference sequences from the CALeDNA Project (Curd et al., 2019). The similarity threshold for CD-HIT is 0.8. Relative purity was calculated by dividing purity by the average of 10 random samples of the same fixed cluster size.

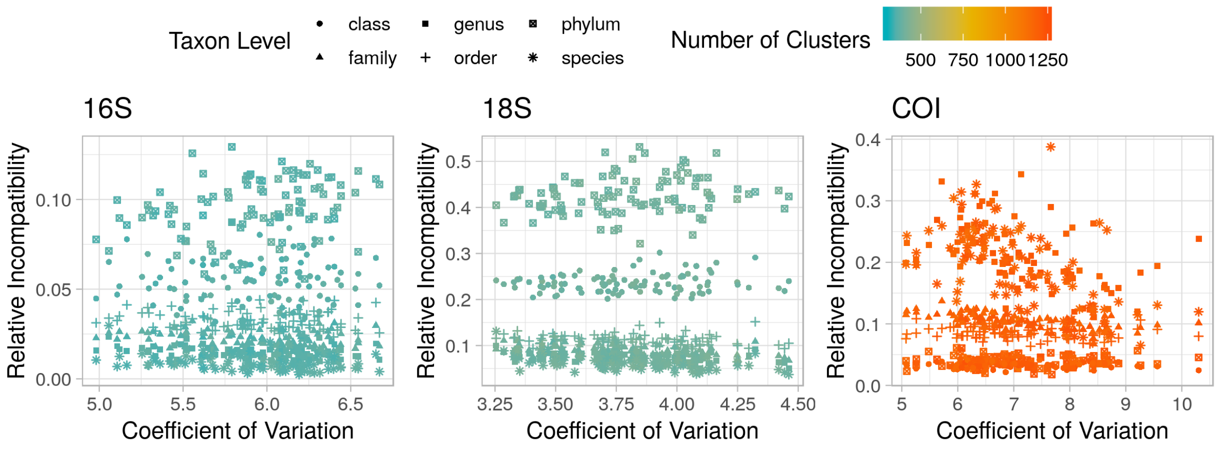


**Figure S8.** Relative incompatibility at all taxon levels for CD-HIT against coefficient of variation for 100 samples of 10,000 randomly chosen 16S, 18S, and COI reference sequences from the CALeDNA Project (Curd et al., 2019). The similarity threshold for CD-HIT is 0.8. Relative incompatibility was calculated by dividing taxonomic incompatibility by the average of 10 random samples of the same fixed cluster size.

Table S1. Taxonomy of sequences used for analysis in Table 1.

| **Taxonomy** |
|---|
| Eukaryota;Apicomplexa;Aconoidasida;Haemosporida;Plasmodiidae;Plasmodium;Plasmodium vivax |
| Eukaryota;Arthropoda;Arachnida;Araneae;Araneidae;Argiope;Argiope bruennichi |
| Eukaryota;Arthropoda;Collembola;NA;Entomobryidae;Entomobrya;Entomobrya sp. BOLD:ACL6239 |
| Eukaryota;Arthropoda;Insecta;Hymenoptera;Diapriidae;NA;Diapriidae sp. BOLD-2016 |
| Eukaryota;Arthropoda;Maxillopoda;Sessilia;Chthamalidae;Notochthamalus;Notochthamalus scabrosus |
| Eukaryota;Chordata;Mammalia;Carnivora;Canidae;Canis;Canis lupus |
| Eukaryota;Echinodermata;Asteroidea;Valvatida;Ophidiasteridae;Linckia;Linckia laevigata |
| Eukaryota;Mollusca;Bivalvia;Mytiloida;Mytilidae;Mytilus;Mytilus trossulus |
| Eukaryota;Mollusca;Gastropoda;NA;Littorinidae;Melarhaphe;Melarhaphe neritoides |
| Eukaryota;Nematoda;Chromadorea;Rhabditida;Strongyloididae;Strongyloides;Strongyloides stercoralis |
| Eukaryota;Platyhelminthes;Cestoda;Diphyllobothriidea;Diphyllobothriidae;Schistocephalus;Schistocephalus solidus |
